## Supplementary figures and images for "Applications of the indole-alkaloid gramine modulate the assembly of individual members of the barley rhizosphere microbiota"

### Figure S1

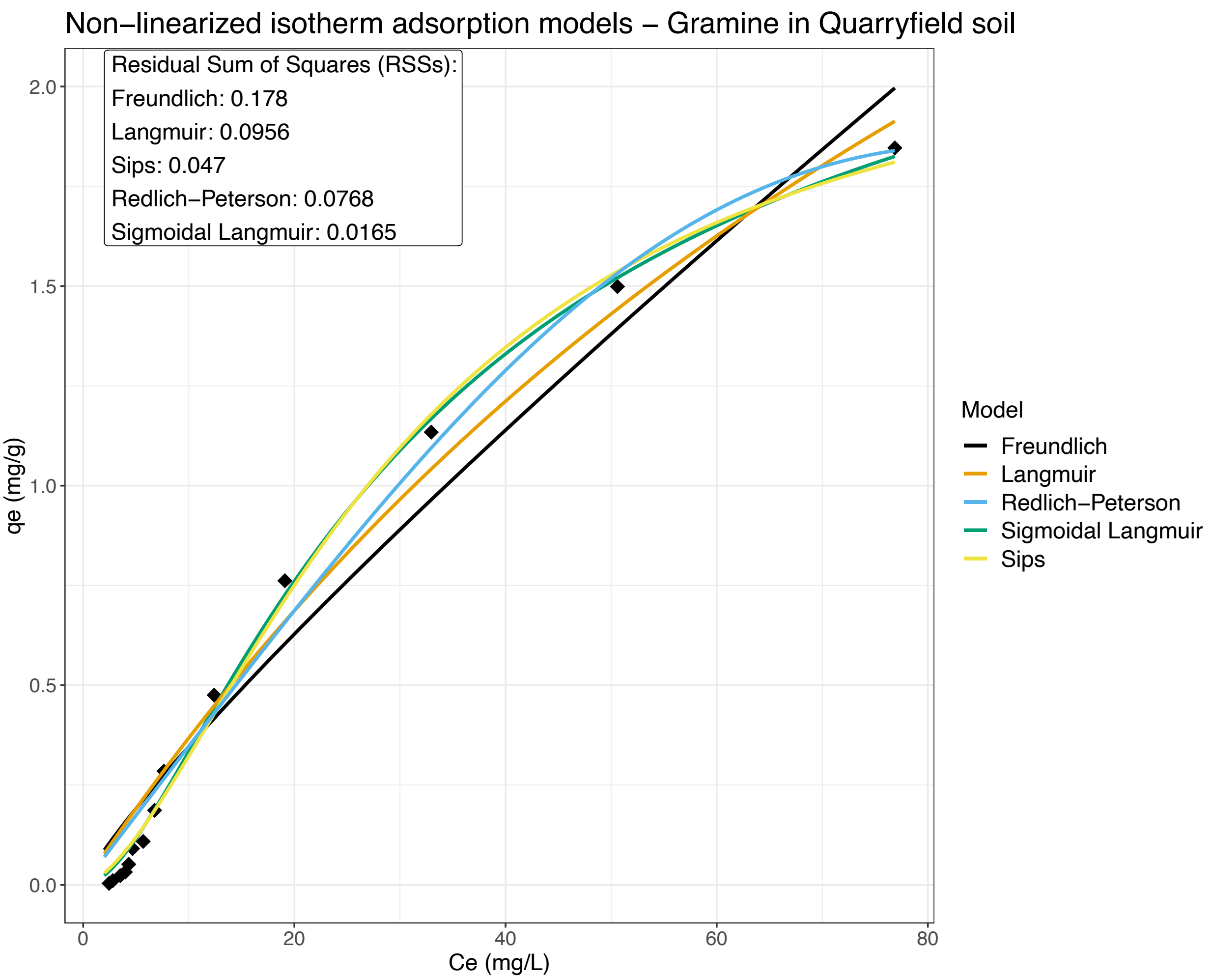

### Figure S2

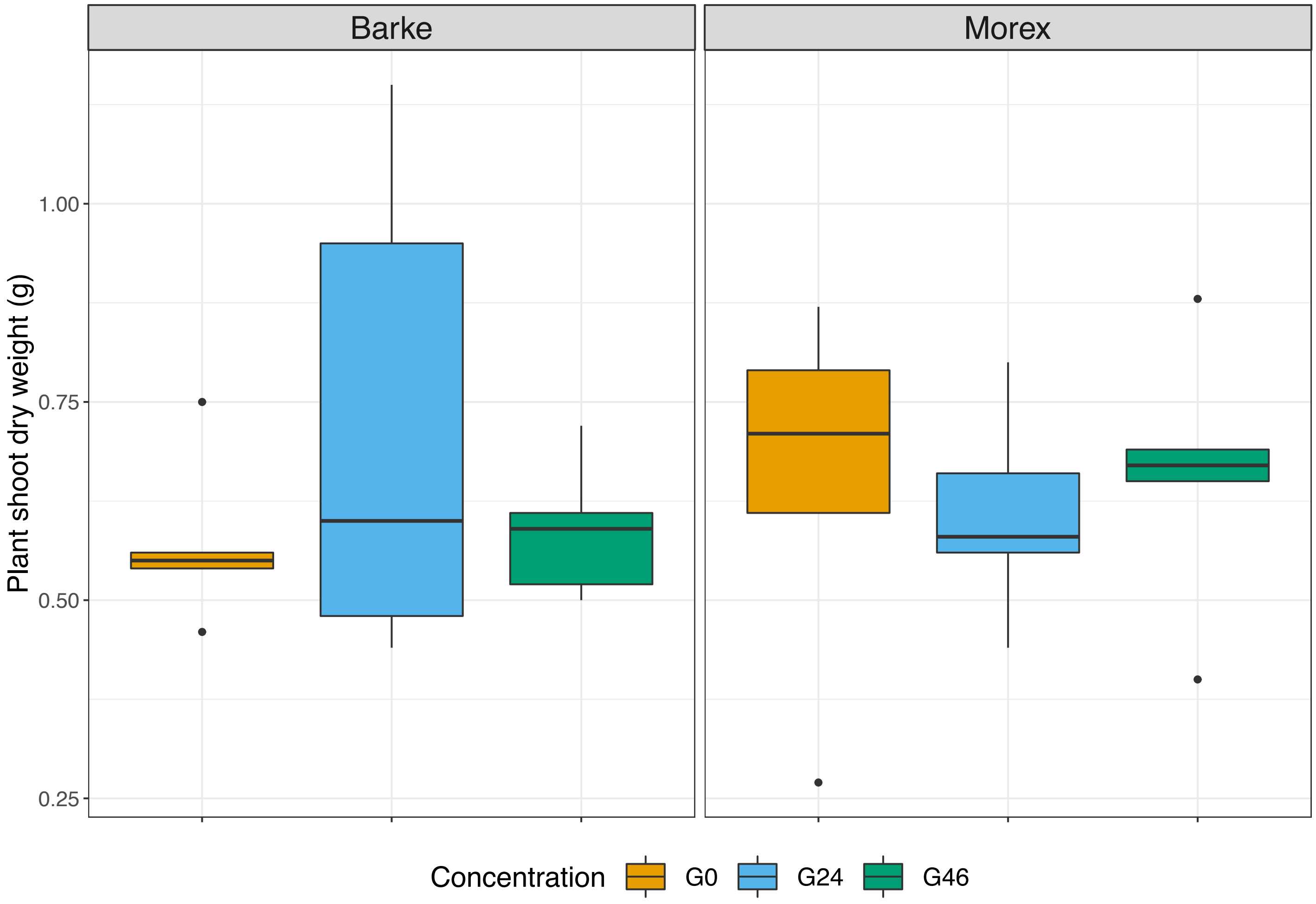

### Figure S3

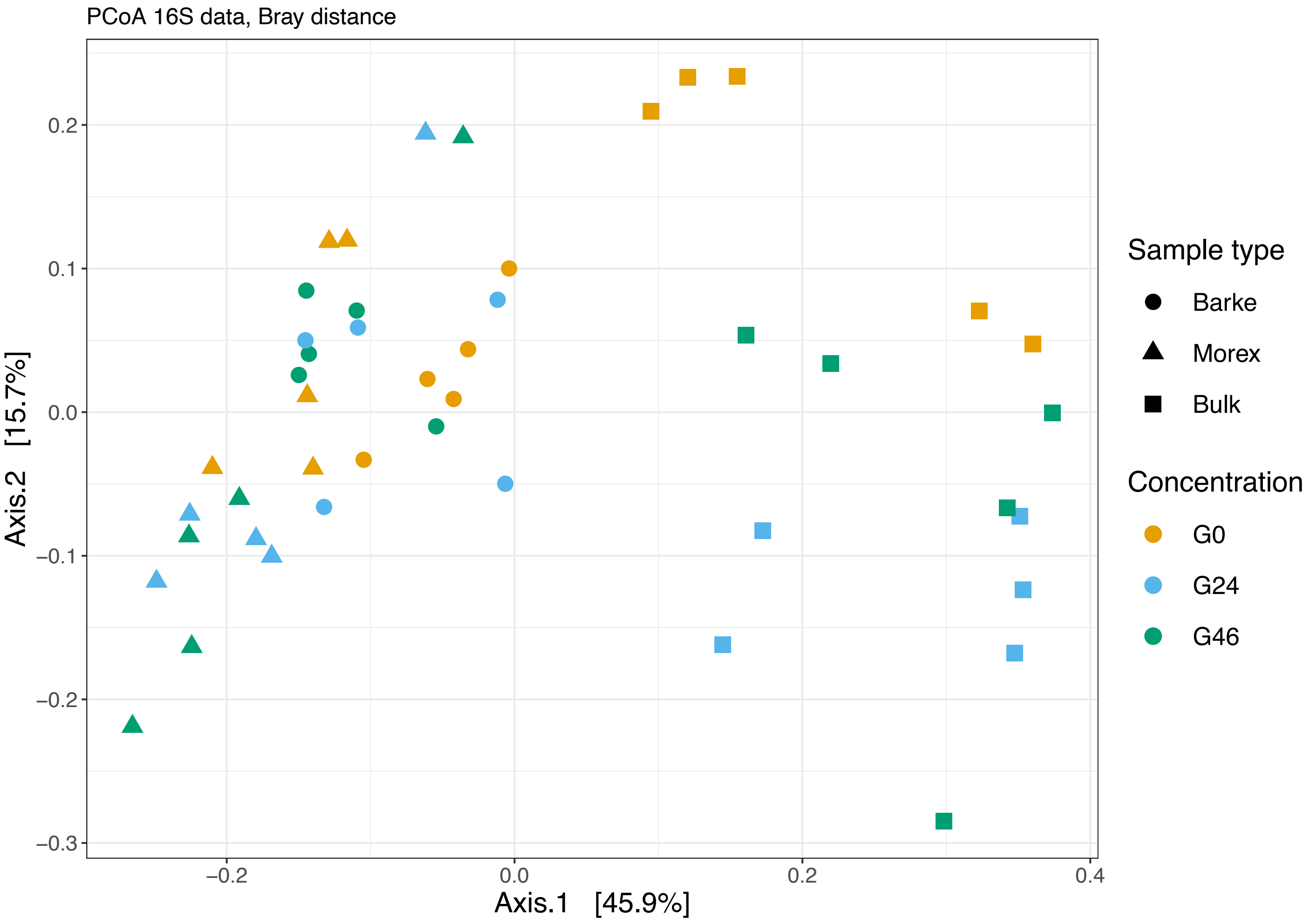

### Figure S4

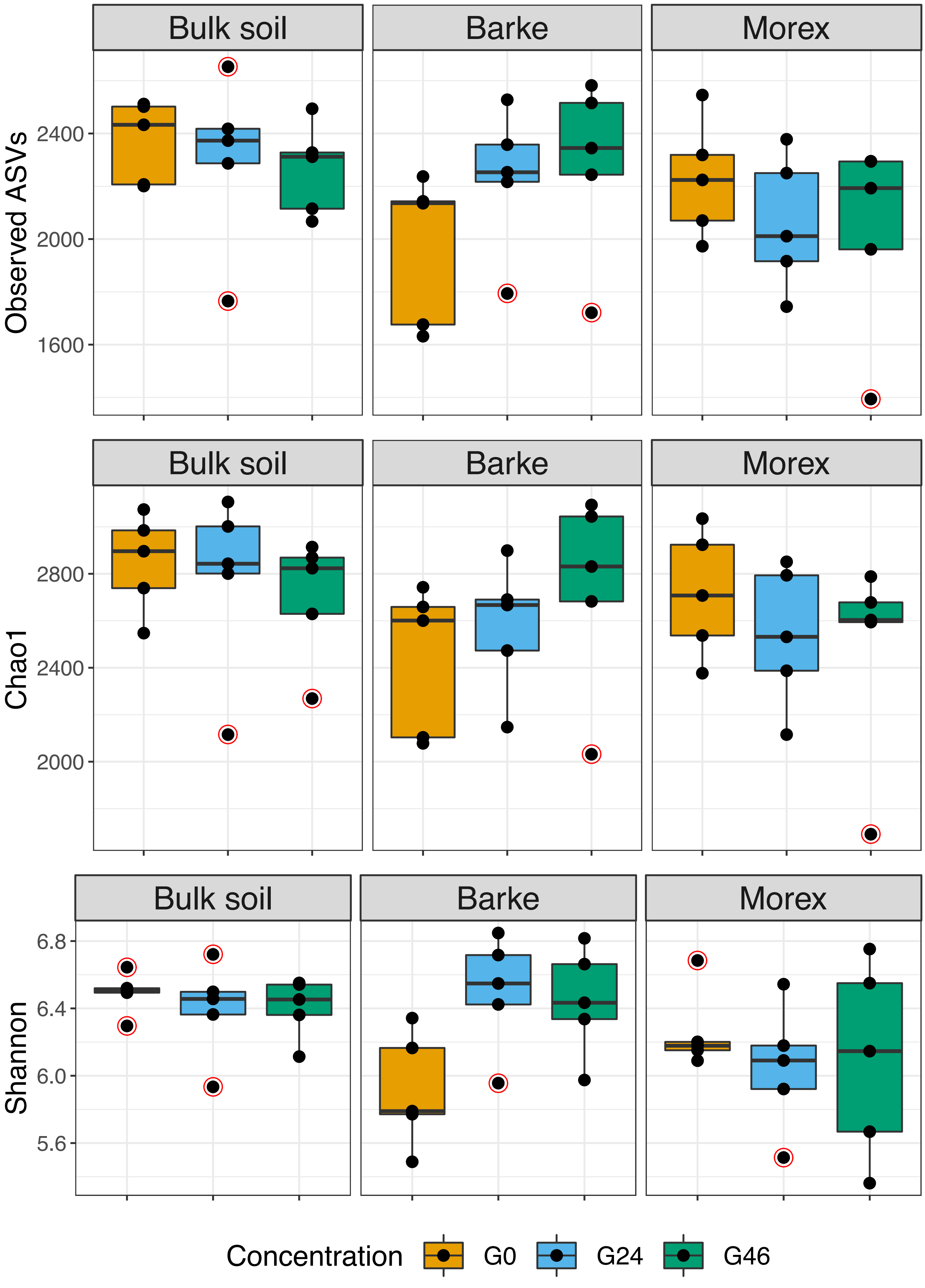

### Figure S5

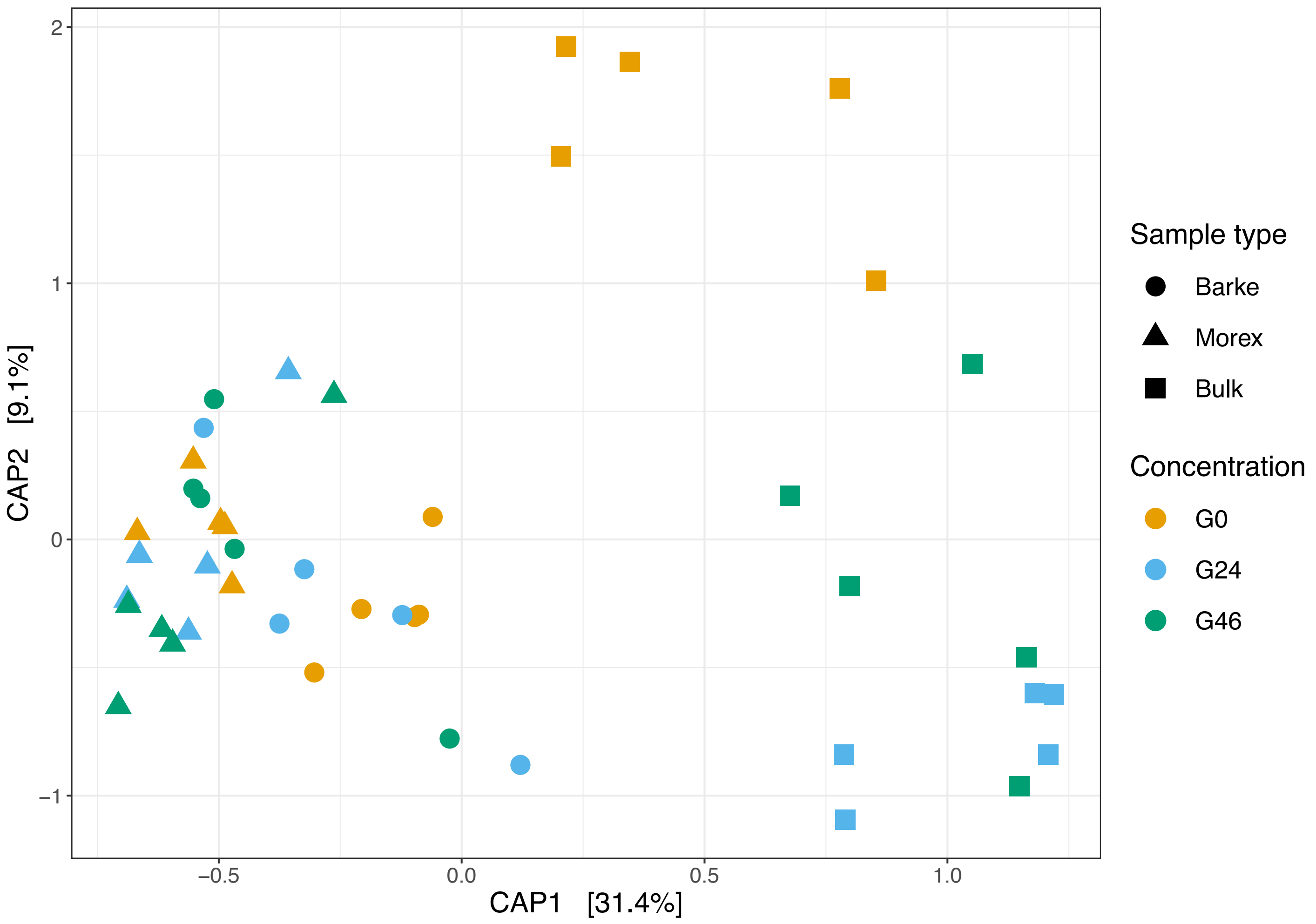
